# AINTEGUMENTA and AINTEGUMENTA-LIKE6 directly regulate floral homeotic and growth genes in young flowers

**DOI:** 10.1101/2020.12.22.424043

**Authors:** Beth A. Krizek, Alexis T. Bantle, Jorman M. Heflin, Han Han, Nowlan H. Freese, Ann E. Loraine

## Abstract

Arabidopsis flower primordia give rise to floral organ primordia in stereotypical positions within four concentric whorls. Floral organ primordia in each whorl undergo distinct developmental programs to become one of four organ types (sepals, petals, stamens, and carpels). The Arabidopsis transcription factors AINTEGUMENTA (ANT) and AINTEGUMENTA-LIKE6 (AIL6) play critical and partially overlapping roles during floral organogenesis. They are required for correct positioning of floral organ initiation, contribute to the specification of floral organ identity, and regulate the growth and morphogenesis of developing floral organs. To gain insight into the molecular means by which ANT and AIL6 contribute to floral organogenesis, we identified the genome-wide binding sites of both ANT and AIL6 in stage 3 flower primordia, the developmental stage at which sepal primordia become visible and class B and C floral homeotic genes are first expressed. AIL6 binds to a subset of ANT sites, suggesting that AIL6 regulates some but not all of the same target genes as ANT. ANT and AIL6 binding sites are associated with genes involved in many biological processes related to meristem and flower organ development. Comparison of genes associated with both ANT and AIL6 ChIP-Seq peaks and those differentially expressed after perturbation of ANT or AIL6 activity identified likely direct targets of ANT and AIL6 regulation. These include the floral homeotic genes *APETALA3* (*AP3*) and *AGAMOUS* (*AG*) and four growth regulatory genes: *BIG BROTHER* (*BB*), *ROTUNDIFOLIA3* (*ROT3*), *ANGUSTIFOLIA3/GRF INTERACTING FACTOR* (*AN3/GIF1*), and *XYLOGLUCAN ENDOTRANSGLUCOLSYLASE/HYDROLASE9* (*XTH9*).

**One Sentence Summary:** The transcription factors ANT and AIL6 directly regulate genes involved in different aspects of flower development including genes that specify floral organ identity and those that regulate growth.

## Introduction

Flowers have long fascinated humans for both their beauty and morphological diversity. In addition, flowers are of practical importance since they contribute to human nutrition in the form of fruits, seeds and grains. In *Arabidopsis thaliana*, flowers arise iteratively from the periphery of the inflorescence meristem, a dome-shaped structure at the apex of the plant. Auxin accumulation within these cells activates MONOPTEROS/AUXIN RESPONSE FACTOR5 (MP/ARF5), a transcription factor that upregulates expression of the floral meristem identity gene *LEAFY* (*LFY*) as well as two *AINTEGUMENTA-LIKE/PLETHORA* (*AIL/PLT*) genes, *AINTEGUMENTA* (*ANT*) and *AIL6/PLT3*, to promote flower primordia identity and outgrowth (Yamaguchi et al., 2013). ANT and AIL6, two members of the larger APETALA2/ETHYLENE RESPONSE FACTOR (AP2/ERF) transcription factor family, and LFY, a plant-specific transcription factor, act early to establish flower primordia and later to promote their continued development with the specification and elaboration of different floral organ types (Weigel et al., 1992; Weigel and Meyerowitz, 1993; Elliott et al., 1996; Klucher et al., 1996; Krizek, 2009; Yamaguchi et al., 2016).

Floral meristems give rise to floral organ primordia at precise positions within four concentric whorls. Floral organ primordia adopt one of four fates (sepal, petal, stamen or carpel) based on their relative position within the developing flower. Positional information within floral primordia is conveyed through distinct combinations of floral organ identity gene activities in each whorl of the flower as summarized in the ABCE model [reviewed in (Wellmer et al., 2014)]. In the outermost whorl one, the class A *[APETALA1 (AP1)* and *APETALA2 (AP2)]* and E genes *[SEPALLATA1-4 (SEP1-4)]* specify sepal identity. In whorl two, the class A, B [*APETALA3* (*AP3*) and *PISTILLATA* (*PI*)] and E genes specify petal identity. In whorl three, class B, C *[AGAMOUS* (*AG*)] and E genes in whorl stamen identity, and in whorl four, class C and E genes specify carpel identity. Mutations in class A, B and C genes results in homeotic changes in floral organ identity in two adjacent whorls of the flower. Thus, these genes are also referred to as floral homeotic genes. All of the class A, B, C, and E genes except the class A gene *AP2* encode MADS domain transcription factors (Yanofsky et al., 1990; Jack et al., 1992; Mandel et al., 1992; Goto and Meyerowitz, 1994; Pelaz et al., 2000; Ditta et al., 2004). AP2 is a member of the AP2/ERF transcription factor family which also includes the AIL/PLT proteins ANT and AIL6 (Jofuku et al., 1994).

Both *lfy* single mutants and *ant ail6* double mutants display loss of floral organ identities but do not show homeotic transformations in organ identity as described for mutations in the class A, B, and C genes (Weigel et al., 1992; Krizek, 2009). *lfy* flowers consists primarily of leaf-like and carpel-like organs and exhibit reduced expression of class B and C genes (Weigel and Meyerowitz, 1993). Broadly-expressed LFY acts with region-specific cofactors to directly activate expression of class B and C genes in stage 3 flowers (Lee et al., 1997; Busch et al., 1999; Lenhard et al., 2001; Lohmann et al., 2001; Lamb et al., 2002; Liu et al., 2009). *ant ail6* double mutants produce flowers with sepals, filaments, stamenoid organs, unfused carpel valves, and structures that do not resemble any wild-type floral organs (Krizek, 2009). *ANT* and *AIL6* act redundantly to promote petal, stamen and carpel identity as these floral organs are present in both *ant* and *ail6* single mutants (Elliott et al., 1996; Klucher et al., 1996; Krizek, 2009). Previous work has shown that expression of the class B genes *AP3, PI* and the class C gene *AG* is reduced in *ant ail6* flowers but it was not known whether ANT and AIL6 directly activate these genes in stage 3 flowers (Krizek, 2009; Krizek et al., 2016).

In addition to defects in floral organ identity, *ant ail6* flowers make fewer and smaller floral organs that do not arise with regular phyllotaxy (Krizek, 2009). *ANT* plays a more important role than *AIL6* in promoting floral organ growth as *ant* floral organs are reduced in size while *ail6* flowers are normal in appearance (Elliott et al., 1996; Klucher et al., 1996; Krizek, 2009). However, the enhanced growth defects in *ant ail6* double mutant flowers indicate that *AIL6* also contributes to organ growth (Krizek, 2009). Overexpression of either *ANT* or *AIL6* can result in larger flowers indicating that both genes are sufficient for floral organ growth (Krizek, 1999; Mizukami and Fischer, 2000; Krizek and Eaddy, 2012; Han and Krizek, 2016).

Here we used chromatin immunoprecipitation in combination with next generation sequencing (ChIP-Seq) to investigate the molecular means by which ANT and AIL6 regulate early events in flower development. We identified genome wide binding sites for both ANT and AIL6 in stage 3 flowers, the stage at which sepal primordia are first visible and class B and C gene expression is initiated. Our work reveals that the partial redundancy of *ANT* and *AIL6* results from AIL6 binding a subset of the genomic regions bound by ANT. Both ANT and AIL6 bind genomic loci associated with genes regulating many different developmental processes including radial patterning, polarity specification, boundary formation, floral meristem determinacy, and floral organ morphogenesis. To identify the mostly likely direct targets of ANT and AIL6 regulation, we compared our ChIP-Seq data with genes previously identified as differentially expressed in either *ant ail6* double mutant inflorescences compared to wild type or those differentially expressed after induction of ANT activity in steroid treated *35S:ANT-GR* inflorescences (Krizek et al., 2016; Krizek et al., 2020). Furthermore, we investigated how the expression of some of these candidate direct target genes respond to inducible downregulation of *AIL6* in the *ant* mutant background. Our data support a direct role for ANT and AIL6 in activation of the floral homeotic genes *AP3* and *AG* in stage 3 flowers and in regulating the expression of four growth regulatory genes, *BIG BROTHER (BB), ROTUNDIFOLIA 3 (ROT3), ANGUSTIFOLIA3/GRF-INTERACTING FACTOR 1 (AN3/GIF1)* and *XYLOGLUCAN ENDOTRANSGLUCOLSYLASE/HYDROLASE9* (*XTH9*).

## Results

### ChIP-Seq identifies many more ANT binding peaks than AIL6 binding peaks

To begin to understand the overlapping functions of ANT and AIL6 in early stages of flower development, we performed ChIP-Seq using previously described ANT-VENUS and AIL6-VENUS fusion lines expressed under their respective endogenous promoters in the *AP1:AP1-GR ap1 cal* synchronized floral induction system (O’Maoiléidigh et al., 2015; Han and Krizek, 2016; Krizek et al., 2020). The *AP1:AP1-GR ap1 cal* system allows for the collection of inflorescences composed of flowers of a single stage of flower development (O’Maoiléidigh et al., 2015).

Inflorescences from *AP1:AP1-GR ap1 cal* (No tag), *ANT:ANT-VENUS ant AP1:AP1-GR ap1 cal*(ANT-VENUS), and *AIL6:AIL6-VENUS AP1:AP1-GR ap1 cal* (AIL6-VENUS) plants were collected two days after dexamethasone (dex) treatment when the inflorescences are composed of stage 3 flowers. Using a visual analytic method in IGB (Integrated Genome Browser (Freese et al., 2016), we identified 2,631 peaks for ANT-VENUS and 595 peaks for AIL6-VENUS. Visual inspection of the data reveals many overlapping ANT and AIL6 peaks. However, there are many more ANT peaks than AIL6 peaks, and ANT peaks are higher in signal than AIL6 peaks. This suggests that ANT regulates more genes than AIL6 and that there is generally lower occupancy of genomic regions by AIL6 as compared with ANT in the collected tissue. This is consistent with the more important role of ANT as compared with AIL6 in floral organ development (Krizek, 2009). Loss of *ANT* function results in smaller floral organs and slight reductions in floral organ number while loss of *AIL6* function has no phenotypic consequence (Elliott et al., 1996; Klucher et al., 1996; Krizek, 2009). The lower signal of the majority of AIL6 binding peaks as compared with corresponding ANT peaks may be a consequence of the lower levels of AIL6 protein as compared with ANT in developing flowers or from AIL6 having lower affinity for these genomic regions as compared with ANT. In addition, competition between the endogenous AIL6 protein and the transgenic AIL6-VENUS protein may have lowered the ChIP-Seq signal in this experiment compared to that for ANT-VENUS. For the ANT-VENUS line, we used an *ant* mutant background in which there is likely no endogenous ANT protein produced.

We used ChIPpeakAnno to identify genes associated with ANT and AIL6 binding peaks (Zhu et al., 2010). ANT peaks were associated with 2,318 unique genes while AIL6 peaks were associated with 592 unique genes (Supplemental Data S1). ANT and AIL6 peaks showed similar distributions relative to the positions of genes. For ANT, half of the peaks (50%) are present upstream of the gene with the remaining peaks either overlapping the start of transcription (18%), within the gene (12%), overlapping the end of transcription (7%), downstream of the gene (12%) or encompassing the gene (1%) (Figure 1A). For AIL6, over half of the peaks (53%) are present upstream of the gene with the remaining peaks either overlapping the start of transcription (18%), within the gene (11%), overlapping the end of transcription (4%), downstream of the gene (13%) or encompassing the gene (1%) (Figure 1C). For both ANT and AIL6, the majority of peaks are located within 2.5 kb of the transcriptional start site (TSS) (Figure 1B, D).

**Figure 1.**
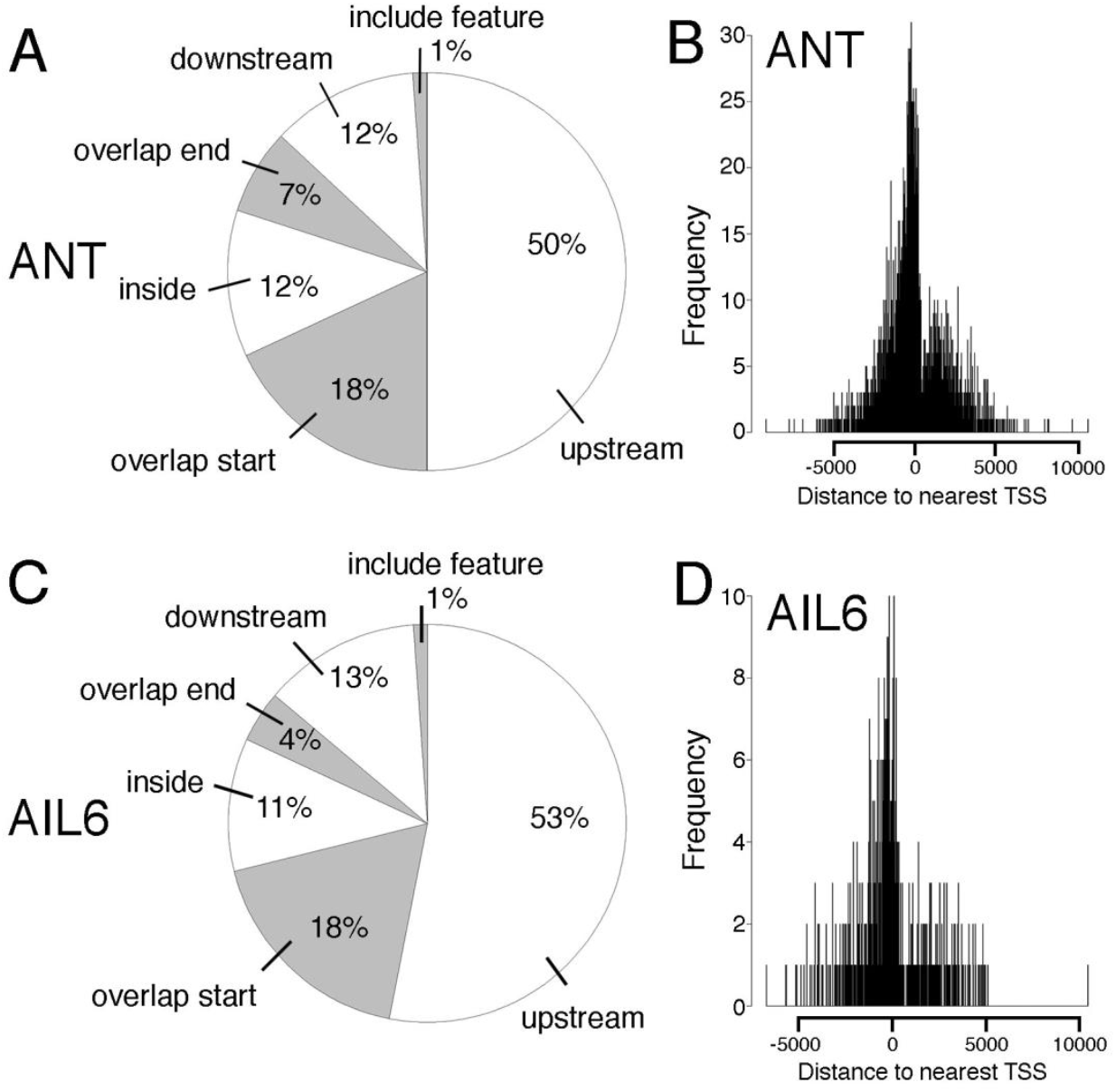
Position of ANT and AIL6 ChIP-Seq peaks relative to closest gene. A. Pie chart showing the position of ANT ChIP-Seq binding peaks relative to the closest gene. Approximately half of the peaks are upstream of the closest gene (50%). The remaining peaks either overlap the start of the gene (18.0%), are within the gene (12%), overlap the end of the gene (7%), are downstream of the gene (12%) or overlap the entire gene (1%). B. Position of ANT binding peaks relative to the transcriptional start site (TSS) of the closest gene. C. Pie chart showing the position of AIL6 ChIP-Seq binding peaks relative to the closest gene. Approximately half of the peaks are upstream of the closest gene (53%). The remaining peaks either overlap the start of the gene (18.0%), are within the gene (11%), overlap the end of the gene (4%), are downstream of the gene (13%) or overlap the entire gene (1%). D. Position of AIL6 binding peaks relative to the transcriptional start site (TSS) of the closest gene.

### GO term enrichment analyses link ANT and AIL6 function with meristem and flower development, hormone physiology and transcriptional regulation

Gene ontology (GO) enrichment analyses identified a number of overrepresented terms in the set of genes associated with either ANT or AIL6 peaks (Figure 2; Supplemental Data S2, S3). Overrepresented GO biological process terms that were identified for both ANT and AIL6 include many associated with meristem and lateral organ development including: polarity specification of adaxial/abaxial axis (GO:0009944), floral meristem determinacy (GO:0010582), radial pattern formation (GO:0009956), cell fate specification (GO:0001708), meristem initiation (GO:0010014), maintenance of meristem identity (GO:0010074), plant ovule development (GO:0048481), and regulation of flower development (GO:0009909) (Figure 2A, B). Other overrepresented biological process development-related GO terms for genes associated with ANT binding peaks include: floral organ formation (GO:0048449), stomatal complex morphogenesis (GO:0010103), anther development (GO:0048653), and leaf morphogenesis (GO:0009965) (Figure 2A). An additional overrepresented GO term for genes associated with AIL6 binding peaks was formation of plant organ boundary (GO:0090691), floral organ morphogenesis (GO:0048444), plant organ formation (GO:1905393) and shoot system morphogenesis (GO:0010016) (Figure 2B).

**Figure 2.**
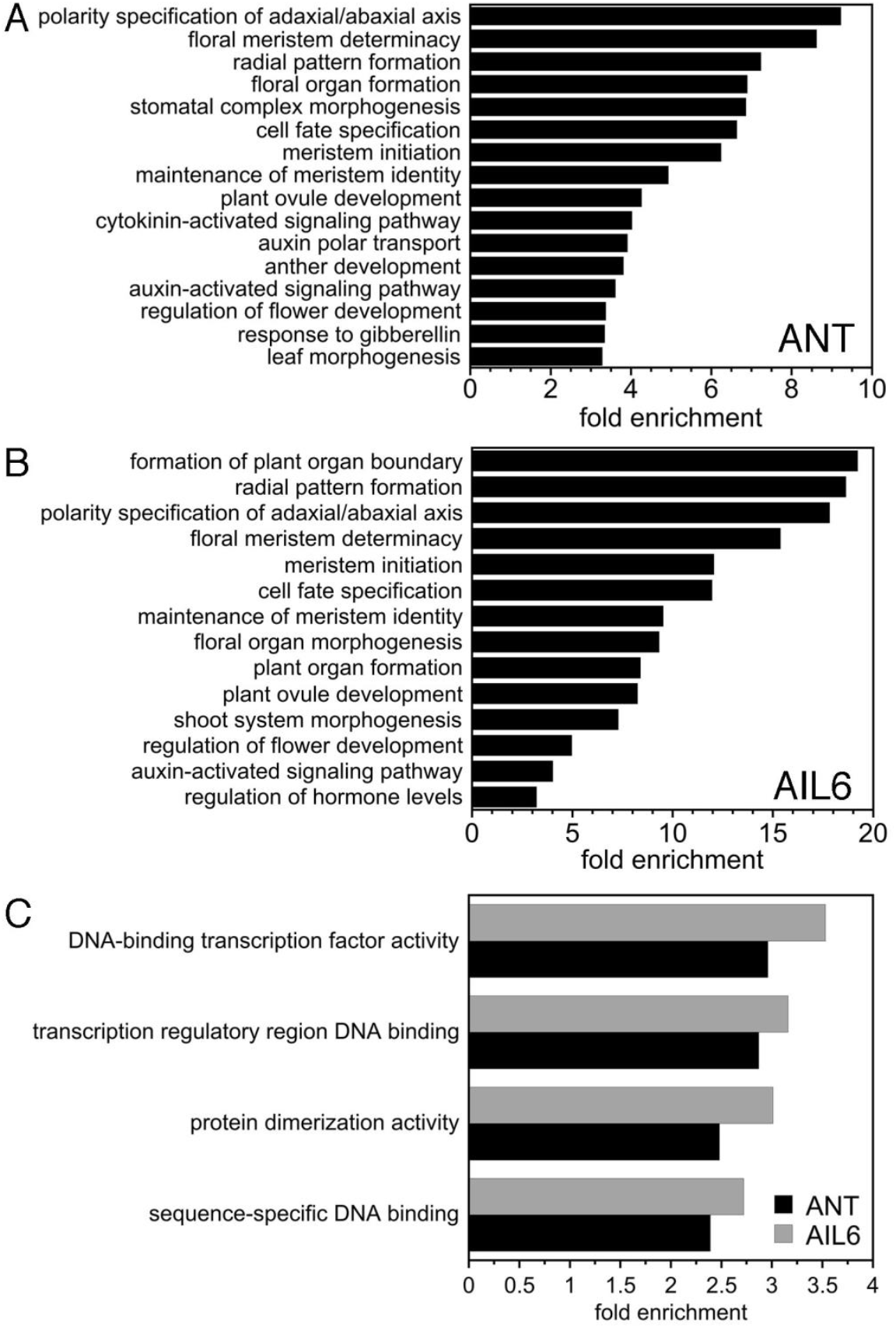
GO enrichment analyses on genes associated with ANT and AIL6 ChIP-Seq binding peaks. A. Biological process GO terms enriched in genes associated with ANT binding peaks. B. Biological process GO terms enriched in genes associated with AIL6 binding peaks. C. Molecular function GO terms enriched in genes associated with ANT and AIL6 binding peaks.

One overrepresented biological process hormone-related GO term associated with both ANT and AIL6 binding peaks was auxin-activated signaling pathway (GO:0009734) (Figure 2A, B). In addition, other overrepresented hormone terms for genes associated with ANT peaks include cytokinin-activated signaling pathway (GO:0009736), auxin polar transport (GO:0009926), response to gibberellin (GO:0009739), ethylene-activated signaling pathway (GO:0009873), and response to abscisic acid (GO:0009737) (Figure 2A, B; Supplemental Data S2). These results suggest that both ANT and AIL6 regulate auxin signaling while ANT plays a more general role in mediating multiple hormone signaling pathways and responses.

Several overrepresented molecular function GO terms for both ANT and AIL6 are associated with transcriptional regulation. These include DNA-binding transcription factor activity (GO:0003700), transcription regulatory region DNA binding (GO:0044212), protein dimerization activity (GO:0046983), and sequence-specific DNA binding (GO:0043565) (Figure 2C; Supplemental Data S2, S3). ANT and AIL6 bind to numerous genomic regions associated with transcription factors that regulate development.

### ANT and AIL6 ChIP-Seq peaks contain DNA binding motifs for AIL/PLT, BPC and bHLH transcription factors

MEME-ChIP from the MEME Suite was used to perform *de novo* motif discovery on the binding peaks for both ANT and AIL6 (Machanick and Bailey, 2011). This analysis used the DAP-Seq database for motif discovery which contains motifs for several AIL/PLT transcription factors including AIL6 but not ANT (O’Malley et al., 2016). AIL/PLT binding motifs are fairly long in length, with a few conserved residues on both ends of the motif and somewhat less conserved nucleotides in the center, as shown for AIL6 (Figure 3). This is less true for the *in vitro* determined ANT binding motif which consists of more conserved residues throughout the motif (Nole-Wilson and Krizek, 2000). For both ANT and AIL6 ChIP-Seq binding peaks, motifs with similarity to those bound by AIL/PLT transcription factors were identified (Table 1). For ANT, these motifs are MEME-2 (NNNNKCMTYGRDWHYYGTG) and DREME-7 (CMTCGRGA), and for AIL6 the single motif MEME-2 (GGCACRHWTYYCRAKGMNN) was identified (Table 1; Figure 3). Both of the MEME motifs contain conserved nucleotides at each end of the AIL/PLT-like binding motif, while the DREME-7 motif identified in ANT binding peaks has similarity to one half of AIL/PLT-like motifs (Figure 3).

**Figure 3.**
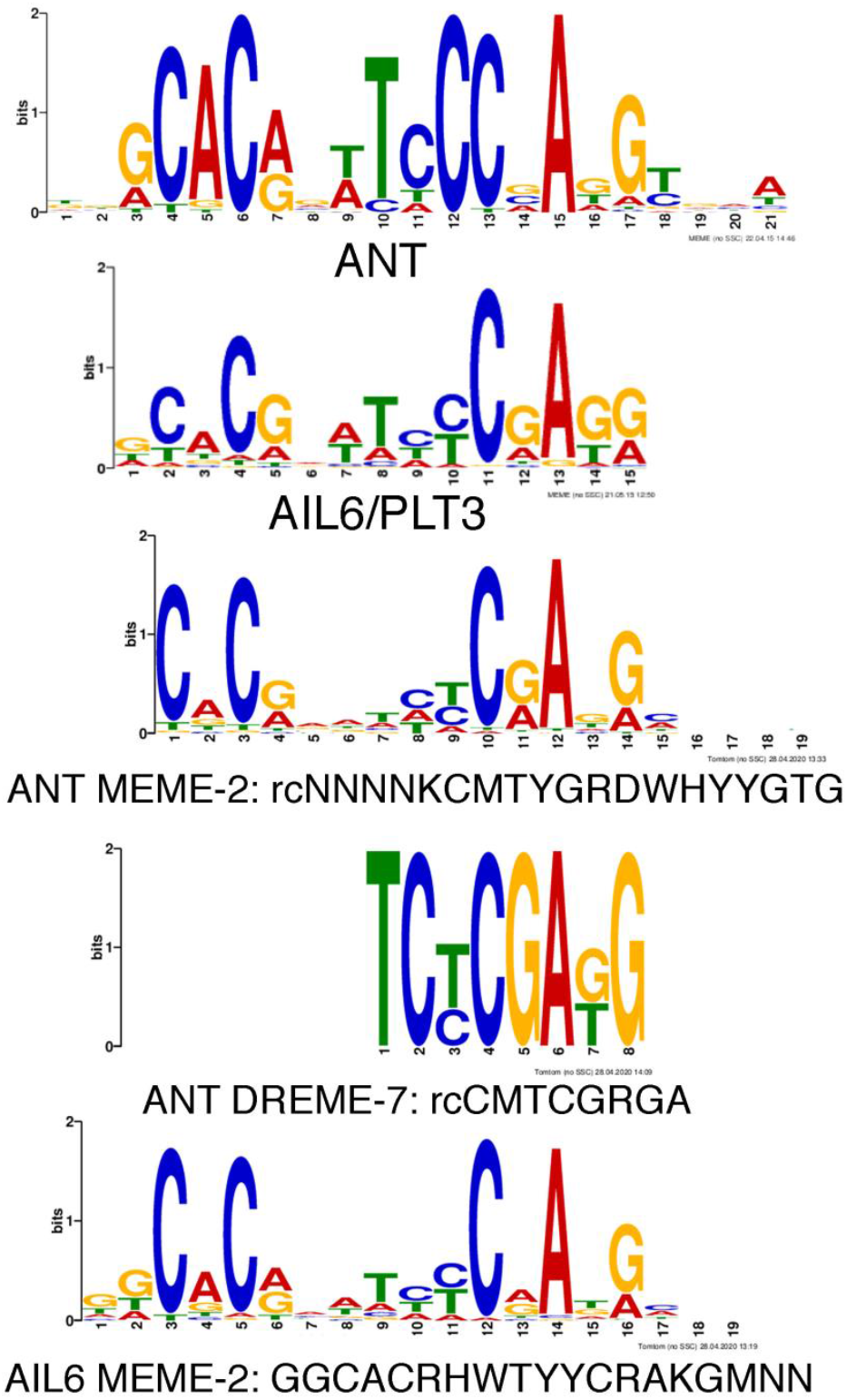
DNA motifs with similarity to ANT and AIL6 binding motifs are overrepresented in ANT and AIL6 ChIP-Seq binding peaks. (Top) Sequence logos for ANT and AIL6 binding motifs. (Middle) Two motifs overrepresented in ANT binding peaks that have similarity to the ANT binding motif are ANT MEME-2 and ANT DREME-7. (Bottom) One motif overrepresented in AIL6 binding peaks that has similarity to the AIL6 binding motif is AIL6 MEME-2. Abbreviation: rc, reverse complement.

**Table 1.**
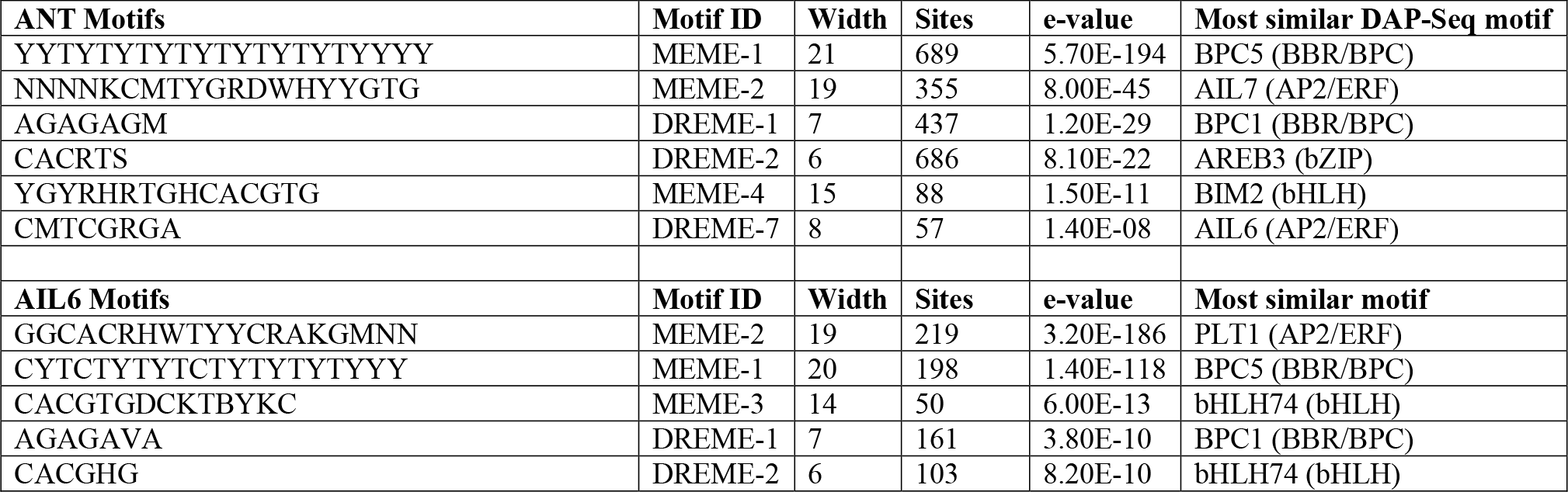
MEME-ChIP analysis of ANT and AIL6 ChIP-Seq peaks

BARLEY B-RECOMBINANT/BASIC PENTACYSTEINE (BBR/BPC) and basic helix loop helix (bHLH) transcription factor binding motifs were overrepresented in both ANT and AIL6 stage 3 binding peaks (Table 1, Supplemental Figure S1). Previously, we found that binding motifs for these two families of transcription factors were overrepresented in ANT binding peaks identified in stage 6 and 7 flowers (Krizek et al., 2020). BBR/BPC transcription factors are broadly expressed transcription factors involved in many developmental processes (Monfared et al., 2011). They mediate gene silencing by recruiting Polycomb repressive complexes or other regulatory proteins to GAGA-motifs (Simonini et al., 2012; Hecker et al., 2015; Xiao et al., 2017). Binding motifs for basic leucine zipper transcription factors (bZIPs) were also overrepresented in ANT binding peaks (Table 1). These other transcription factors may act in combination with ANT and AIL6 in regulating transcription of target genes.

### AIL6 binding peaks correspond to a subset of ANT peaks

We determined the degree of overlap between ANT and AIL6 binding peaks. 582 out of the 595 AIL6 peaks (approximately 98%) overlap at least 50% with an ANT peak (Figure 4A). For the 13 AIL6 peaks that did not overlap at least 50% with an ANT peak, this was because there was no nearby ANT peak (three), the AIL6 peak was wider with a summit shifted compared with an ANT peak in the same region (three), or there was a overlapping an ANT peak but its height was below the 2.5 threshold value used for peak identification (seven) (Supplemental Data S4). These results indicate that AIL6 binds almost exclusively to a subset of regions also bound by ANT in stage 3 flowers. This is consistent with AIL6 providing some but not all of the same functions as ANT (Krizek, 2009). 606 unique genes are associated with the 582 overlapping peaks between ANT and AIL6 (Supplemental Data S5). GO enrichment analyses on this set of genes suggest that ANT and AIL6 have overlapping roles in regulating plant organ boundaries, radial pattern formation, polarity specification, and floral organ morphogenesis (Figure 4B; Supplemental Data S6). Some of the genes bound by both ANT and AIL6 in these GO term categories are shown in Table 2.

**Figure 4.**
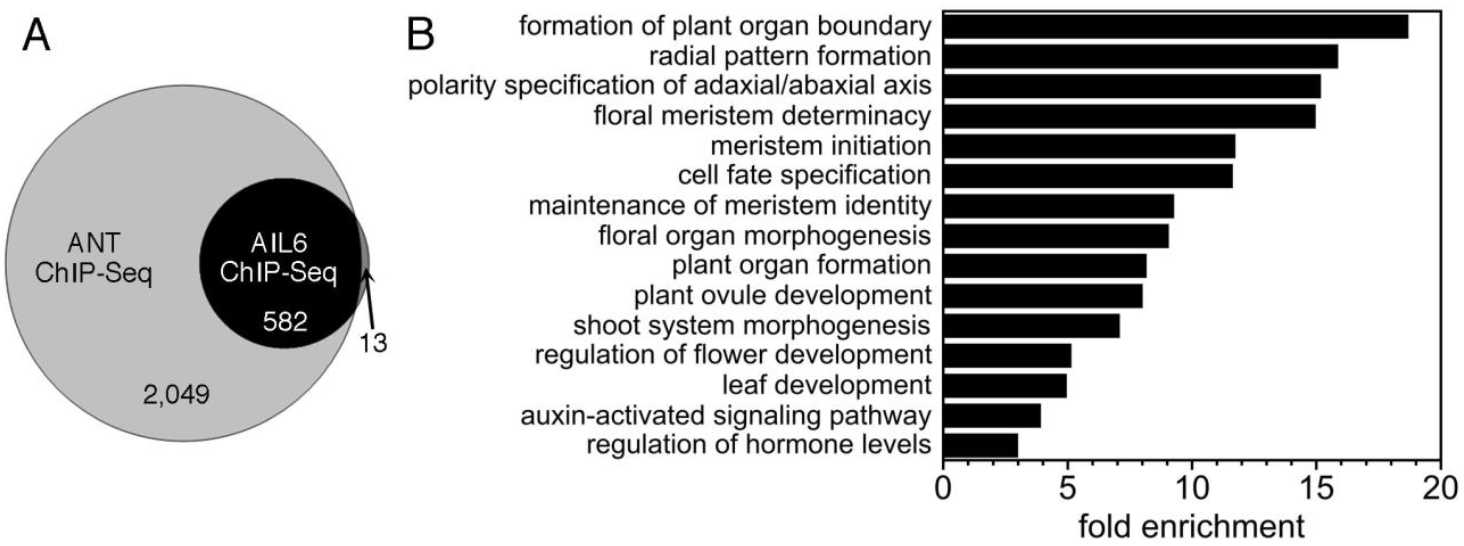
Overlap of ANT and AIL6 ChIP-Seq data. A. Venn diagram showing the overlap of ANT and AIL6 ChIP-Seq binding peaks. B. Biological process GO terms enriched in genes associated with both ANT and AIL6 binding peaks.

**Table 2.**
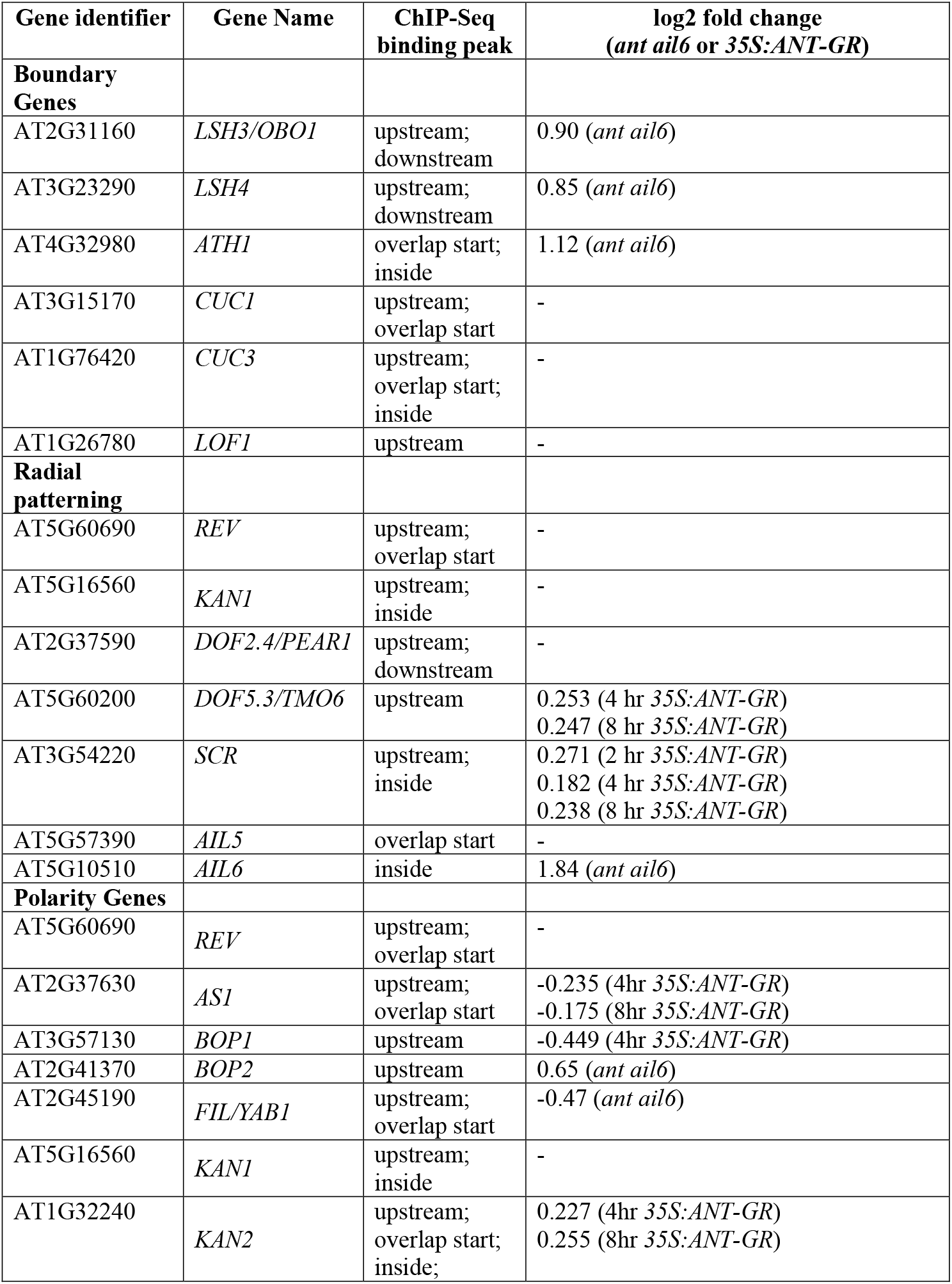
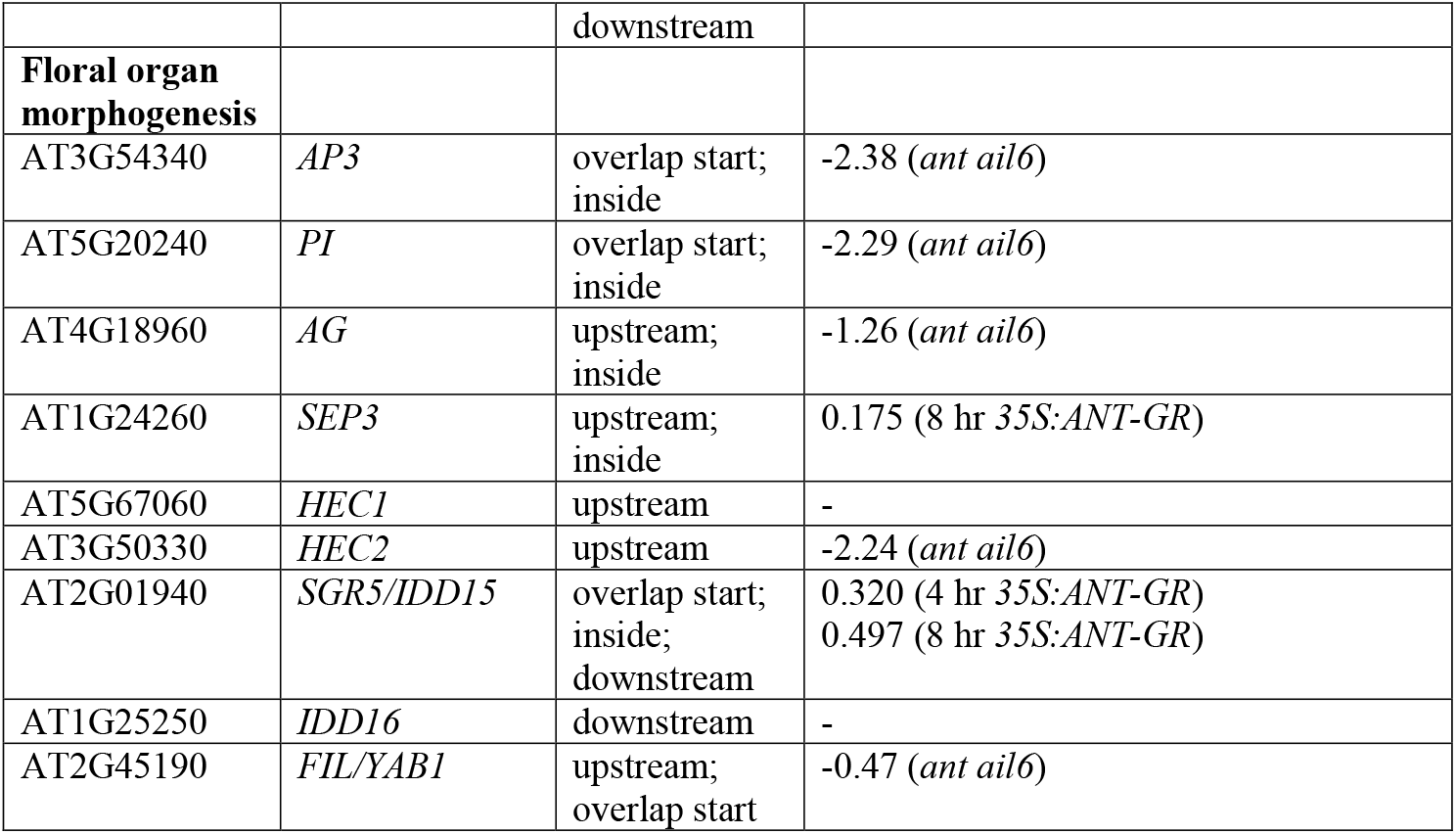
Developmental genes associated with both ANT and AIL6 ChIP-Seq peaks

### 175 genes bound by ANT and AIL6 are differentially expressed in *ant ail6*

Direct targets of a transcription factor are typically defined as genes whose regulatory region is bound by the transcription factor and whose expression is altered in response to changes in the activity of the transcription factor. While ChIP-Seq can identify many hundreds or thousands of transcription factor binding sites, not all of the genes associated with these sites may be transcriptionally regulated by these binding events. Jointly analyzing ChIP-Seq and RNA-Seq offers the best approach to identify direct target genes. A comparison of the set of genes bound in stage 3 flowers by both ANT and AIL6 (606 genes) with the set of genes that are differentially expressed in *ant ail6* inflorescences (8,012 genes) identified an overlap of 175 genes (Figure 5A; Table 2) (Krizek et al., 2016). GO ontology enrichment analyses on these genes identifies the following biological process GO terms: maintenance of meristem identity (GO:0010074), plant organ formation (GO:1905393), floral organ development (GO:0048437), post-embryonic plant morphogenesis (GO:0090698), regionalization (GO:0003002), phyllome development (GO:0048827), and response to hormone (GO:0009725) (Figure 5C; Supplemental Data S7). In addition, this analysis identified the following overrepresented molecular process GO terms: protein dimerization activity (GO:0046983), DNA-binding transcription factor activity (GO:0003700), and sequence-specific DNA binding (GO:0043565), and the cellular component GO term glyoxysome (GO:0009514) (Supplemental Data S7).

**Figure 5.**
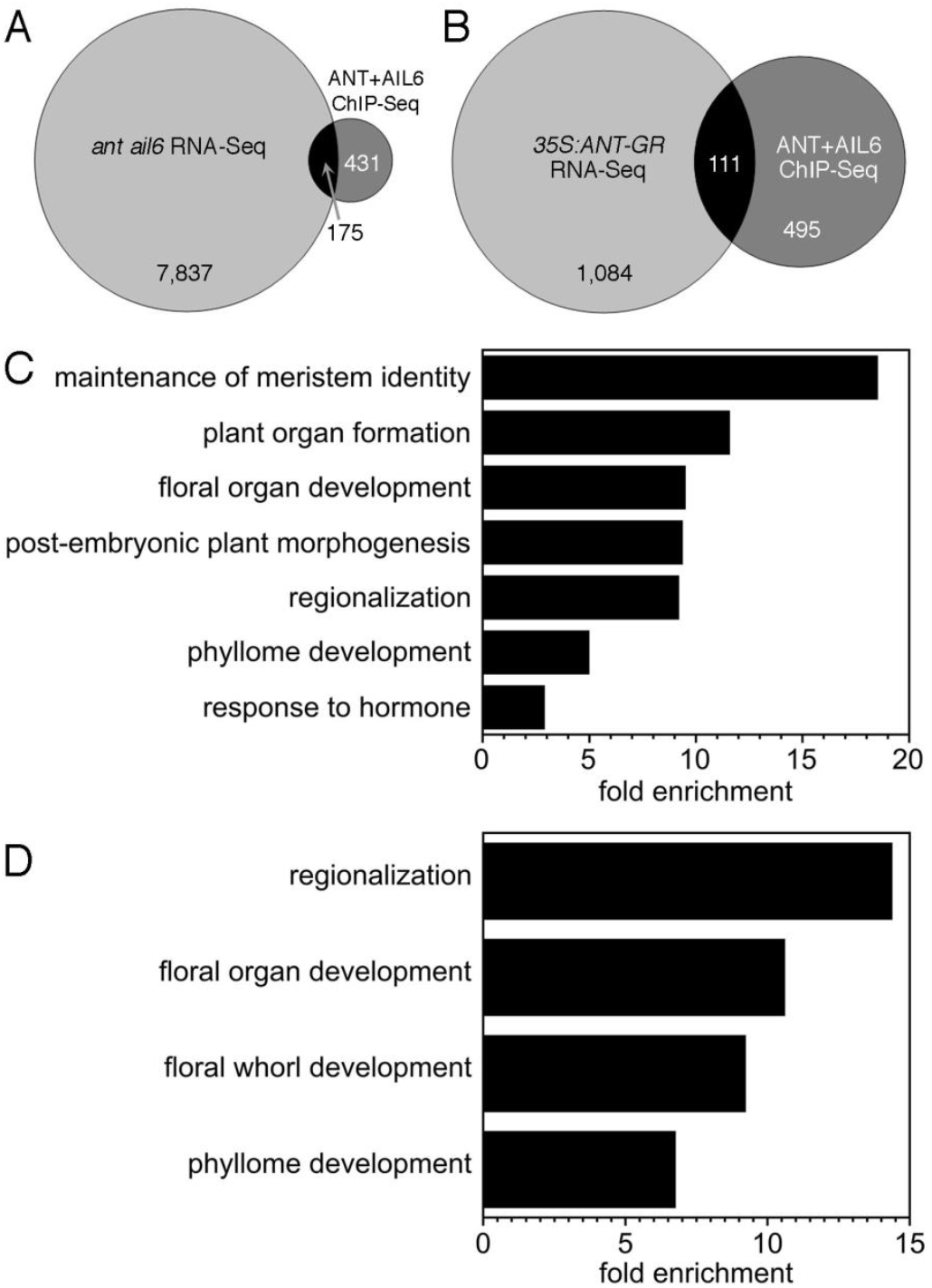
Venn diagrams and GO enrichment analyses on set of genes that are bound by both ANT and AIL6 and differentially expressed in *ant ail6* and *35S:ANT-GR* RNA-Seq experiments. A. Venn diagram showing the overlap of genes associated with both ANT and AIL6 ChIP-Seq peaks and genes differentially expressed in *ant ail6* compared to *Ler* RNA-Seq experiment. B. Venn diagram showing the overlap of genes associated with both ANT and AIL6 ChIP-Seq peaks and genes differentially expressed in dex treated *35S:ANT-GR* compared to mock treated *35S:ANT-GR* RNA-Seq experiment. C. Biological process GO terms enriched in genes associated with both ANT and AIL6 binding peaks and differentially expressed in *ant ail6* inflorescences. D. Biological process GO terms enriched in genes associated with both ANT and AIL6 binding peaks and differentially expressed in *35S:ANT-GR* inflorescences after dex treatment.

### 111 genes bound by ANT and AIL6 are differentially expressed *35S:ANT-GR*

We performed a second comparison of the genes bound in stage 3 flowers by both ANT and AIL6 (606 genes) with genes that are differentially expressed after induction of ANT activity in *35S:ANT-GR* inflorescences (1,195 genes) (Krizek et al., 2020). This comparison identified 111 genes that are likely direct targets of ANT and AIL6 regulation (Figure 5B; Table 2). GO ontology enrichment analyses on these genes identifies the following biological process GO terms: regionalization (GO:0003002), floral organ development (GO:0048437), floral whorl development (GO:0048438), and phyllome development (GO:0048827) (Figure 5D; Supplemental Data S8). In addition, this analysis identified the following molecular function GO term: DNA-binding transcription factor activity (GO:0003700) (Supplemental Data S8).

### ANT and AIL6 activate class B and C homeotic genes in stage three flowers

Included in the set of 175 genes that are both bound by ANT and AIL6 and differentially expressed in *ant ail6* inflorescences are the class B floral homeotic genes *AP3* and *PI*, and the class C floral homeotic gene *AG* (Figure 6A, D; Figure S2A, Table 2). We confirmed the ChIP-Seq results for *AP3, PI*, and *AG* using ChIP-qPCR (Figure 6B, C, E, F; Figure S2B, C). Expression of these three genes is downregulated in *ant ail6* inflorescences (Krizek et al., 2016). In addition, in situ hybridization had shown previously that *AP3* and *AG* were expressed in fewer cells of stage 3 floral primordia (Krizek, 2009). These results are consistent with the loss of petal, stamen and carpel identities in *ant ail6* flowers (Krizek, 2009). These data suggest that ANT and AIL6 might directly activate the expression of class B and C floral homeotic genes in young flowers.

**Figure 6.**
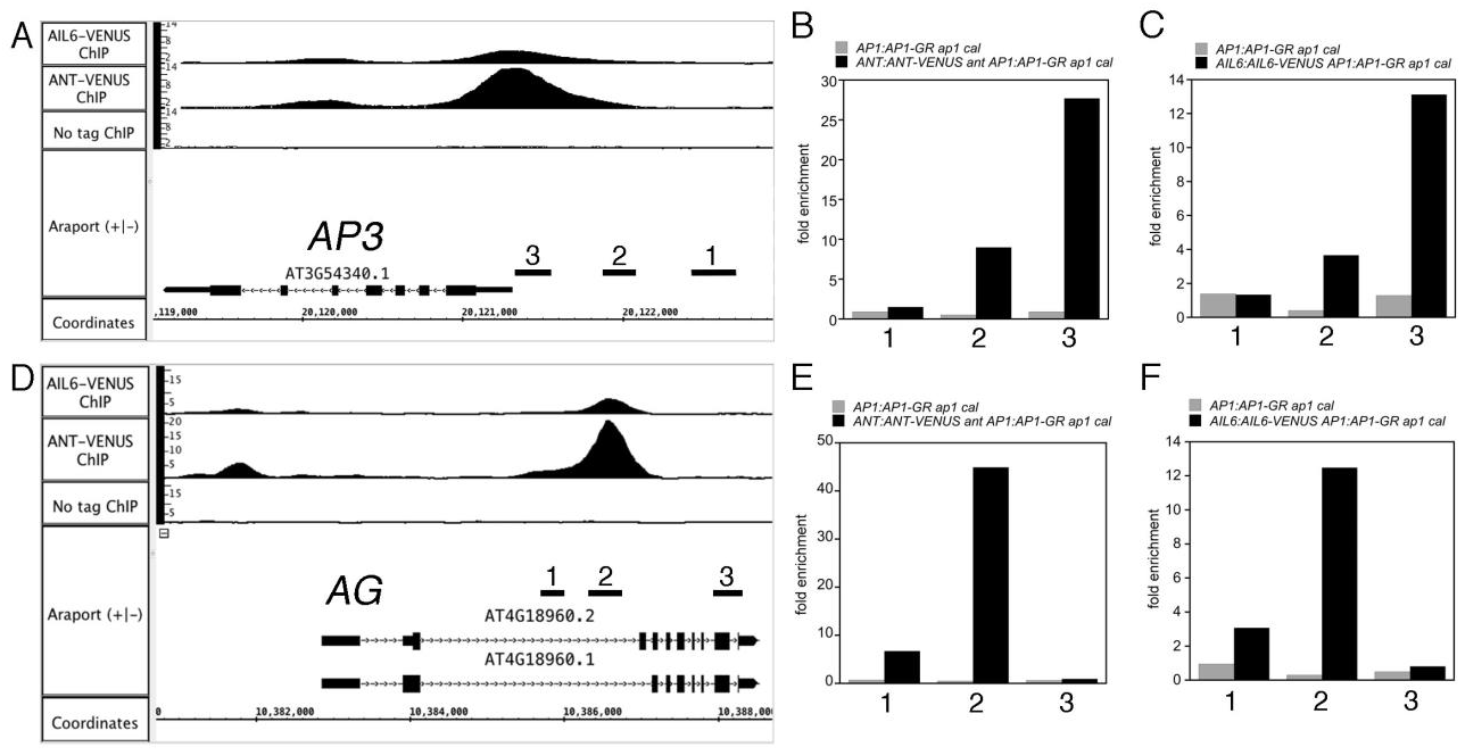
ANT and AIL6 bind directly regulatory regions associated with the floral organ identity genes *AP3* and *AG.* A. ChIP-Seq of ANT and AIL6 binding to *AP3* genomic region. 1, 2, and 3 are genomic regions tested for binding using ChIP-qPCR in B and C. B. ChIP-qPCR of ANT binding to *AP3* genomic region 3. C. ChIP-qPCR of AIL6 binding to *AP3* genomic region 3. D. ChIP-Seq of ANT and AIL6 binding to *AG* genomic region. 1, 2, and 3 are genomic regions tested for binding using ChIP-qPCR in E and F. E. ChIP-qPCR of ANT binding to *AG* genomic region 2. F. ChIP-qPCR of AIL6 binding to *AG* genomic region 2.

To investigate whether ANT and AIL6 directly control transcription of *AP3, PI* and *AG*, we examined whether class B and C gene expression responds quickly to changes in ANT and AIL6 activity. We used an ethanol inducible transgenic line in which *AIL6* expression is downregulated by an artificial microRNA (amiR) in the *ant* mutant background *(35S:ALCR/AlcA:AIL6-amiR ant;* i.e. *AIL6-amiR ant).* After an eight-hour ethanol treatment, *AIL6-amiR ant* flowers exhibit a phenotype more severe than *ant* with loss of petals and partially unfused carpels, suggesting that AIL6 activity is reduced (Figure 7A). Mock treated *AIL6-amiR ant* flowers resemble *ant* (Figure 7A). *AIL6* mRNA levels in the ethanol treated inflorescences are approximately 30% of those in mock treated inflorescences, indicating that the *AIL6-amiR* knocks down *AIL6* expression (Figure 7B). *AP3* and *AG* mRNA levels but not *PI* mRNA levels are reduced after this eight-hour ethanol treatment (Figure 7B; Figure S2D). These data further support the conclusion that ANT and AIL6 directly activate expression of the class B gene *AP3* and the class C gene *AG* in stage 3 flowers.

**Figure 7.**
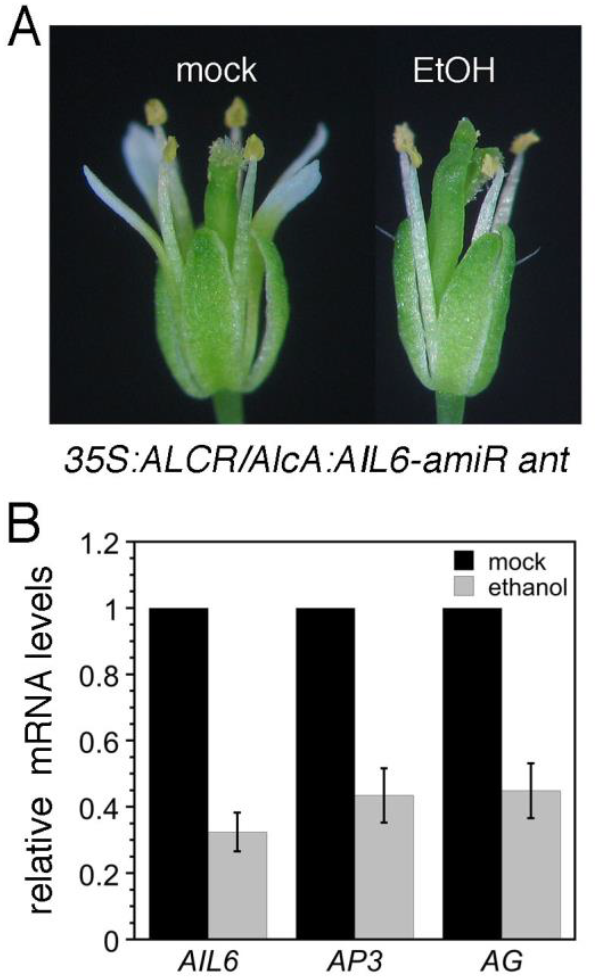
*AP3* and *AG* expression is reduced soon after downregulation of *AIL6* expression in *35S:ALCR/AlcA:AIL6-amiR ant* inflorescences. A. Mock and ethanol (EtOH) treated *35S:ALCR/AlcA:AIL6-amiR ant* flowers. B. Expression of *AIL6, AP3* and *AG* is reduced in *35S:ALCR/AlcA:AIL6-amiR ant* inflorescences after an eight-hour EtOH treatment.

No DNA sequences with obvious similarity to ANT or AIL6 binding motifs are present in *AP3* genomic regions (Nole-Wilson and Krizek, 2000; O’Malley et al., 2016; Krizek et al., 2020). The ANT MEME-2 and AIL6 MEME-2 motifs identified here were also not present in these peaks. Thus, is it not clear what DNA sequence is bound by ANT and AIL6 within the *AP3* genomic region. The ANT and AIL6 binding peaks upstream of *AP3* do overlap the defined PEE core element that is required for early stage 3-5 expression (Lamb et al., 2002). Two partially overlapping sequences with weak similarity to the ANT binding motif are present near the summit of the ANT and AIL6 ChIP-Seq binding peaks within the *AG* intron (Supplemental Figure S3A). ANT can bind specifically to this region of the *AG* intron *in vitro* and can activate transcription in yeast through this sequence (Supplemental Figure S3B,C). For both *AP3* and *AG*, the ANT and AIL6 binding peaks overlap those of other known regulators of *AP3, PI*, and *AG*, including LFY, AP1, AP2, AP3, PI, AG and SEP3 (Supplemental Figure S4A,B).

### Cross-regulation of *AIL* expression may involve direct repression by ANT and AIL6

Previously, we found evidence of cross-regulation among *AIL* gene expression (Krizek et al., 2016). Specifically, we found that *AIL6* mRNA levels are increased in *ant* mutants and that *ANT, AIL6* and *AIL7* mRNA levels are increased in *ant ail6* double mutants (Krizek et al., 2016). Our ChIP-Seq results show that both ANT and AIL6 bind the regulatory regions of *ANT, AIL5*, and *AIL6;* ANT also binds to the regulatory region of *AIL7* (Supplemental Figure S5). This suggests that the observed repression of *AIL* expression by ANT and AIL6 may be mediated by direct binding of ANT and AIL6 to *AIL* regulatory regions.

As seen with the floral homeotic genes, ANT and AIL6 binding peaks in these regulatory regions overlap those of other floral regulators, including LFY, AP1, AP3, PI, and AG (Supplemental Figure S5). *ANT* and *AIL6* expression in floral anlagen occurs with similar timing as *LFY* expression and LFY was previously shown to bind to *ANT* and *AIL6* genomic regions (Winter et al., 2011). Thus, LFY may contribute to *ANT* and *AIL6* expression in floral primordia. In contrast, *AP1*, *AP3, PI* and *AG* are expressed later in flower development than *ANT* and *AIL6* and thus would only contribute to maintenance or refinement of *ANT* and *AIL6* expression at later stages of development, if they play a role at all.

### ANT and AIL6 directly regulate organ growth genes

*ANT* and *AIL6* both positively contribute to floral organ growth, although *AIL6* cannot completely substitute for *ANT* in this role as *ant* single mutants produce smaller floral organs (Elliott et al., 1996; Klucher et al., 1996). Several growth-regulating genes are bound by both ANT and AIL6 in our ChIP-Seq experiments and exhibit differential expression in either *ant ail6* double mutants or dex treated *35S:ANT-GR* inflorescences. These include the growth repressor *BIG BROTHER (BB)*, and the growth-promoting genes *ROTUNDIFOLIA3 (ROT3), ANGUSTIFOLIA3/GRF-INTERACTING FACTOR 1 (AN3/GIF1)* and *XYLOGLUCAN ENDOTRANSGLUCOSYLASE/HYDROLASE9 (XTH9)* (Supplemental Figure S6). *BB* expression was elevated in *ant ail6* double mutants, *ROT3* expression was reduced in *ant ail6* double mutants, and *AN3/GIF1* and *XTH9* expression was upregulated by induction of ANT-GR activity (Krizek et al., 2016; Krizek et al., 2020). *AN3/GIF1* and *XTH9* were previously identified as likely direct targets of ANT regulation as their regulatory regions are bound by ANT in stage 6/7 flowers (Krizek et al., 2020).

We further investigated whether these genes are likely to be direct targets of ANT and AIL6 regulation by examining their expression in response to downregulation of *AIL6* in the *ant* mutant background and downregulation of *ANT* alone. To downregulate *ANT* expression, we generated an RNAi line that utilizes an ethanol-inducible inverted repeat transgene *(35S:ALCR/AlcA:ANT-IR;* i.e. *ANT-IR).* After ethanol treatment, *ANT-IR* flowers produce smaller petals and anthers with two locules similar to *ant* mutants (Figure 8A-C). After a 24-hour ethanol treatment, *ANT* mRNA levels are reduced to approximately 25% of the mock treated plants (Figure 8D). *BB* expression was increased 1.8-fold in ethanol-treated *ANT-IR* and 3.1-fold in ethanol-treated *AIL6-amiR ant* inflorescences suggesting that both ANT and AIL6 contribute to repression of *BB* expression (Figure 8E). After ethanol induction, *ROT3* expression levels were 63% of the levels seen in mock treated *ANT-IR* and 42% of the levels seen in mock treated *AIL6-amiR* inflorescences (Figure 8F). *AN3/GIF1* expression was slightly reduced in *ANT-IR* inflorescences and reduced approximately 2-fold in *AIL6-amiR* inflorescences (Figure 8G). This suggests that for *AN3/GIF1* regulation, AIL6 can largely compensate for loss of ANT. After ethanol induction, *XTH9* expression was reduced to approximately 3-fold in both *ANT-IR* and *AIL6-amiRNA ant* inflorescences (Figure 8H).

**Figure 8.**
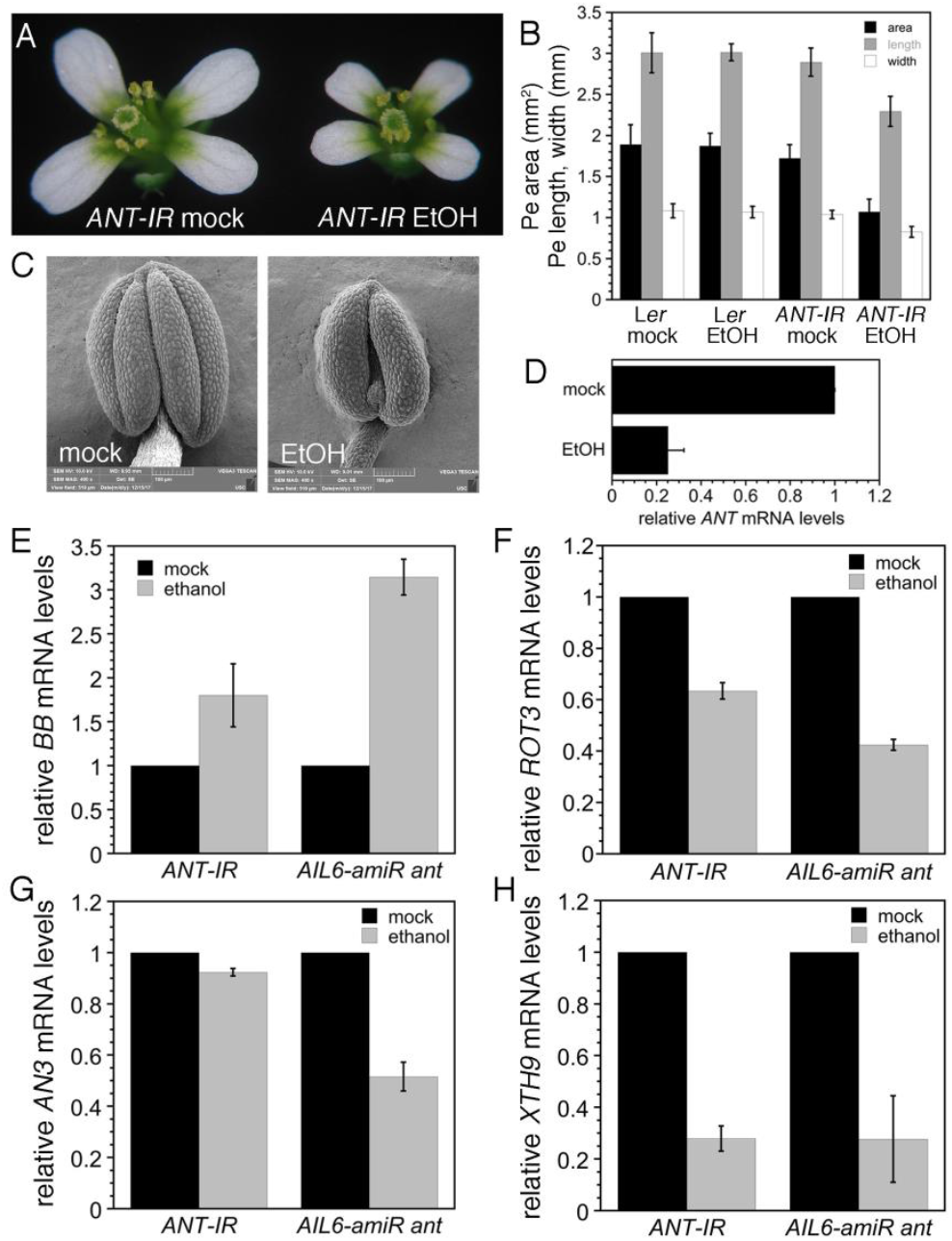
Expression of growth regulatory genes is altered in response to downregulation of *ANT* expression in *35S:ALCR/AlcA:ANT-IR* inflorescences and downregulation of *AIL6* expression in *35S:ALCR/AlcA:AIL6-amiR ant* inflorescences. A. Mock and ethanol (EtOH) treated *35S:ALCR/AlcA:ANT-IR* flowers. B. Petal area, length and width measurements for mock and EtOH treated *Ler* and *35S:ALCR/AlcA:ANT-IR* flowers. C. Stamen anther of mock treated *35S:ALCR/AlcA:ANT-IR* flower. D. Stamen anther of EtOH treated *35S:ALCR/AlcA:ANT-IR* flower. E. Expression of *ANT* is reduced in *35S:ALCR/AlcA:ANT-IR* inflorescences after a 24hour EtOH treatment. F. *BB* expression is upregulated in *35S:ALCR/AlcA:ANT-IR* and *35S:ALCR/AlcA:AIL6-amiR ant* inflorescences after a 24-hour EtOH treatment. G. *AN3* expression is downregulated in *35S:ALCR/AlcA:AIL6-amiR ant* inflorescences after a 24-hour EtOH treatment. *AN3* expression is slightly downregulated in *35S:ALCR/AlcA:ANT-IR* inflorescences after a 24-hour EtOH treatment. H. *XTH9* expression is downregulated in *35S:ALCR/AlcA:ANT-IR* and *35S:ALCR/AlcA:AIL6-amiR ant* inflorescences after a 24-hour EtOH treatment.

## Discussion

Our data indicate that the partially overlapping roles of ANT and AIL6 in flower development are a consequence of these transcription factors regulating many of the same target genes. The more important role of ANT in floral organogenesis as compared with AIL6 appears to result from ANT regulating additional genes that are not targets of AIL6 regulation. In addition, at most of these genomic sites, ANT exhibits higher occupancy than AIL6 which may lead to a more significant effect on transcriptional regulation at these sites. Higher occupancy by ANT may be a consequence of greater amounts of ANT protein as compared with AIL6 rather than intrinsic differences in the DNA binding affinities of ANT and AIL6, as previous work has shown that AIL6 can compensate for loss of ANT function when AIL6 is expressed under the control of the *ANT* promoter (Han and Krizek, 2016).

GO analyses on genes associated with ANT and AIL6 ChIP-Seq peaks suggest that these transcription factors regulate a number of different processes during early stages of flower development. In particular, terms associated with floral meristem development (meristem initiation, maintenance of meristem identity and floral meristem determinacy) and floral organ development (polarity specification, formation of plant organ boundary, radial pattern formation, and floral organ morphogenesis) were identified. Our work here suggests previously unknown roles for ANT and AIL6 in boundary specification and radial pattern formation. Some of the genes bound by both ANT and AIL6 in these processes are also differentially expressed in either *ant ail6* inflorescences or after steroid induction of ANT activity (Table 1). Polarity specification genes had been previously identified as overrepresented in ANT ChIP-Seq binding sites in stage 6/7 flowers and a role for ANT in polarity specification had been suggested by genetic work (Nole-Wilson and Krizek, 2006; Krizek et al., 2020). Our work here also links AIL6 to polarity specification.

We show that ANT and AIL6 directly activate the class B gene *AP3* and the class C gene *AG* in stage 3 flowers. ANT and AIL6 bind to *AP3* and *AG* regulatory regions, expression of *AP3* and *AG* is reduced soon after downregulation of AIL6 activity in the *ant* mutant background, and *AP3* and *AG* are expressed in fewer cells in stage 3 flowers (Krizek, 2009). The regulation of *AG* expression by ANT appears to be complex as *ANT* acts with *APETALA2 (AP2)* to repress *AG* expression in second whorl cells, although it is not known if this regulation is direct (Krizek et al., 2000). Thus, ANT and AIL6 may directly activate *AG* expression in third and fourth whorl cells while repressing (directly or indirectly) *AG* expression in second whorl cells. How broadly-expressed ANT and AIL6 activate *AP3* specifically in second and third whorl cells and *AG*specifically in third and fourth whorl cells is not clear. It is possible that ANT and AIL6 act with region-specific cofactors as has been shown previously for LFY (Lee et al., 1997; Busch et al., 1999; Lenhard et al., 2001; Lohmann et al., 2001; Lamb et al., 2002; Liu et al., 2009). Another question still to be answered is the DNA sequence bound by ANT and AIL6 within the *AP3*promoter. No obvious AIL/PLT-like binding motif was identified within the ChIP-Seq binding peaks. Two overlapping sequences with weak similarity to the ANT binding motif are present within the *AG* intron and bound by ANT *in vitro*. This weak similarity suggests that there may be flexibility in the DNA sequences recognized by ANT. These sequences are located near the LFY and WUSCHEL (WUS) binding sites within the *AG* intron (Busch et al., 1999; Lohmann et al., 2001). The ANT and AIL6 ChIP-Seq binding peaks within both *AP3* and *AG* overlap with those of other floral regulatory genes (Supplemental Figure S4). Thus, these genomic regions appear to contain binding sites for multiple transcription factors that may act cooperatively to regulate transcription. In particular, ANT and AIL6 may act in combination with other transcriptional regulators like LFY, SEP3, AP1, and AP2 to control the spatial and temporal expression of *AP3* and *AG*.

ANT and AIL6 contribute to other aspects of floral organogenesis including growth; this appears to involve the regulation of both growth promoting and growth repressing genes. ANT and AIL6 directly bind to a region near the TSS of the growth repressor *BB*, and *BB* expression is increased in *ant ail6* double mutant inflorescences (Krizek et al., 2016). *bb* mutants produce larger flowers than wild type while overexpression of *BB* results in smaller flowers (Disch et al., 2006). *BB* is expressed throughout young stage 1-4 flowers, similar to *ANT* expression in young flowers and overlapping with *AIL6* (Elliott et al., 1996; Disch et al., 2006; Krizek, 2009). Previous work has shown that transgenic *bb* plants containing a genomic copy of *BB* with a deletion in an upstream region corresponding to the location of the ANT and AIL6 ChIP-Seq binding peaks produced smaller flowers than wild type. The reduced size of these flowers suggesting that these transgenic plants have higher levels of *BB* expression than wild type and that this region contains a negative cis-regulatory element bound by ANT and AIL6 that represses transcription of *BB*. BB is an E3 ubiquitin ligase that ubiquitinates and activates the protease DA1 and is subsequently cleaved by DA1 (Disch et al., 2006; Dong et al., 2017). DA1 limits the duration of the cell proliferation phase of organ growth by cleaving the deubiquitylase UBP15 and two TEOSINTE BRANCHED 1/ CYCLOIDEA/PCF transcription factors, TCP15 and TCP22 (Dong et al., 2017). While BB protein levels are regulated by DA1, ANT and AIL6 are the first identified potential regulators of *BB* transcription.

ANT and AIL6 also bind to regions associated with three growth-promoting genes, *ROT3*, *AN3* and *XTH9.* Previously we showed that ANT binds to the regulatory regions of *AN3* and *XTH9* in stage 6/7 flowers and that both of these genes are activated upon dex induction of ANT-GR activity (Krizek et al., 2020). Here we show that ANT and AIL6 bind to these genomic regions in stage 3 flowers. Furthermore, reduced expression of *ROT3, AN3* and *XTH9* upon downregulation of *AIL6* in the *ant* mutant background provides additional evidence that ANT and AIL6 directly regulate the expression of these genes. This suggests that ANT regulates a variety of growth regulating pathways as ROT3 is a cytochrome P450 acting in brassinosteroid biosynthesis, AN3 is a transcriptional co-activator that works with GROWTH REGULATING FACTOR (GRF) transcription factors, and XTH enzymes modify xyloglucan in the cell wall (Kim et al., 1998; Rose et al., 2002; Kim and Kende, 2004).

In summary, our work provides new insights into the roles of ANT and AIL6 in the initiation and development of floral organs from the floral meristem. We have identified direct targets of ANT and AIL6 regulation that mediate their roles in the establishment floral organ identity and the promotion of floral organ growth. Further work is needed to determine whether other genes identified here are direct targets of these transcription factors that may mediate additional roles for ANT and AIL6 in processes such as boundary formation and radial patterning.

## Materials and Methods

### Plant materials and growth conditions

*ANT:ANT-VENUS ant-4 AP1:AP1-GR ap1 cal* plants and *AIL6:AIL6-VENUS* plants were described previously (Han and Krizek, 2016; Krizek et al., 2020). *AIL6:AIL6-VENUS* plants were crossed to *AP1:AP1-GR ap1cal* plants. *AP1:AP1-GR ap1 cal, ANT:ANT-VENUS ant-4 AP1:AP1-GR ap1 cal*, and *AIL6:AIL6-VENUS AP1:AP1-GR ap1 cal* inflorescences were grown on a soil mixture of Fafard 4P:perlite:vermiculite (8:1:1) in 24 hour days at a light intensity of approximately 160 micromol/m^2^/s at 20°C. An artificial microRNA (amiRNA) that targets AIL6 was designed using http://wmd3.weigelworld.org/cgi-bin/webapp.cgi. A gene fragment containing this AIL6-amiR within the miR319a precursor was synthesized by IDT and cloned into the EcoRI/BamHI sites of BJ36_AlcA to create AlcA:AIL6-amiR/BJ36. AlcA:AIL6-amiR was subcloned into the NotI site of pMLBart_AlcR to create 35S:ALCR/AlcA:AIL6-amiR/pMLBart. 35S:ALCR/AlcA:AIL6-amiR/pMLBart was transformed into Agrobacterium strain ASE, which was then used to transform *ant-4/+* plants. *35S:ALCR/AlcA:AIL6-amiR* transgenic plants were selected for Basta resistance. The first 736 nucleotides of the *ANT* coding region were cloned in the sense and antisense directions in pHannibal using ANTIR-5 and ANTIR-6 (Table S1). The ANT-IR fragment was digested from pHannibal with XhoI/BamHI and cloned into BJ36_AlcA to create AlcA:ANT-IR. AlcA:ANT-IR was subcloned into the NotI site of pMLBart_AlcR to create 35S:ALCR/AlcA:ANT-IR/pMLBart. 35S:ALCR/AlcA:ANT-IR/pMLBart was transformed into Agrobacterium strain ASE, which was then used to transform L*er* plants. *35S:ALCR/AlcA:ANT-IR* transgenic plants were selected for Basta resistance. *35S:ALCR/AlcA:AIL6-amiR ant-4* and *35S:ALCR/AlcA:ANT-IR* plants were grown on a soil mixture of Fafard 4P:perlite:vermiculite (8:1:1) in 16 hour days at a light intensity of approximately 160 micromol/m^2^/s at 22°C.

### ChIP-Seq and ChIP-qPCR

Plants for ChIP-Seq and ChIP-qPCR were treated by pipetting a dex (10μM dexamethasone +0.015% Silwet) solution onto the inflorescences. Chromatin immunoprecipitation was performed as described previously except that inflorescences were collected two days after dex treatment when the tissue consists of stage 3 flowers (Krizek et al., 2020). Primers for ChIP-qPCR are listed in Table S1. Fold enrichment was determined relative to a negative control, the transposon *TA3.* Sequencing libraries were prepared from two biological replicates of input and ChIP DNA for *AP1:AP1-GR ap1 cal, ANT:ANT-VENUS ant-4 AP1:AP1-GR ap1 cal*, and *AIL6:AIL6-VENUS AP1:AP1-GR ap1 cal* as described previously (Krizek et al., 2020). Sequencing was performed on an Illumina HiSeq 2500 producing 150 base paired-end reads. Sequence reads were aligned to the reference *Arabidopsis thaliana* genome (version TAIR9, released June 2009) using bowtie2. Examination of the coverage graphs revealed high reproducibility between the two ChIP-Seq replicates. In addition, the input samples closely resembled the control untagged *AP1:AP1-GR ap1 cal* samples. ANT and AIL6 binding peaks were identified using a visual analytics approach within the Integrated Genome Brower (IGB) (Freese et al., 2016). Specifically, coverage graphs were generated for the combined data from the two replicates and normalized. A difference coverage graph was generated by subtracting coverage graphs of the untagged sample *(AP1:AP1-GR ap1 cal)* from the coverage graphs for the tagged samples *(ANT:ANT-VENUS ant-4 AP1:AP1-GR ap1 cal* and *AIL6:AIL6-VENUS AP1:AP1-GR ap1 cal).* Peaks were defined using the thresholding feature. A thresholding value of 2.5 identified 595 peaks for AIL6 and 2,631 peaks for ANT in stage 3 flowers. For each identified peak, ChIPpeakAnno was used to identify the gene with the closest transcription start site (TSS) (Zhu et al., 2010). In some cases, a peak located within a gene is associated with two genes if the TSS of an adjacent gene is closer than the TSS of the gene overlapping the peak. Gene Ontology analyses were performed with AmiGO 2 (http://amigo.geneontology.org/amigo). BEDtools intersect was used to identify overlapping ANT and AIL6 peaks in stage 3 flowers. *De novo* motif discovery was performed with MEME-ChIP. Source code for bioinformatic analyses are available in the project “git” repository https://bitbucket.org/krizeklab/antail6stage3chipseq/. Venn diagrams were created using Venn Diagram Plotter (omics.pnl.gov) created by the Pacific Northwest National Lab (PNNL).

### RT-qPCR

*35S:AlcR/AlcA:AIL6-amiR ant-4* and *35S:ALCR/AlcA:ANT-IR* plants were treated with mock or ethanol vapor by placing 2mls of water or 2mls of 100% ethanol in 2ml centrifuge tubes in half of the pots in a tray. The tray was covered with a plastic dome. Inflorescences were collected at the end of an eight-hour *(35S:AlcR/AlcA:AIL6-amiR ant-4)* or 24-hour *(35S:ALCR/AlcA:ANT-IR)* treatment. RNA was isolated using an RNeasy Plant Mini Kit (Qiagen) or TRIzol (Life Technologies). Samples isolated with TRIzol were further purified on an RNeasy column (Qiagen) and treated with DNase while on the column. First strand complementary DNA (cDNA) synthesis was performed using Quanta qScript cDNA SuperMix (Quanta BioSciences) following the manufacturer’s instructions. qPCR was performed on a BioRad CFX96 using PerfeCTa SYBR Green FastMix for iQ (Quanta BioSciences) and primers listed in Table S1. Data analyses were carried out as described previously (Krizek and Eaddy, 2012). At least two biological replicates were analyzed for each experiment.

### Gel mobility shift assays

The gel mobility shift assays were performed as described previously (Nole-Wilson and Krizek, 2000).

### Yeast strain and ß-galactosidase assays

A reporter plasmid was made by cloning the 76bp *AG* intron sequence upstream of the *lacZ* gene in the vector pLacZi (Clontech). The yeast reporter strain was made by integration of the linearized *AG* intron reporter plasmid into the yeast strain YM4271. This new reporter strain was transformed with the previously described ANT/pGAD424 lacking the GAL4 activation domain (Krizek, 2003). Transformants were selected on plates containing synthetic medium lacking leucine. ß-galactosidase assays were performed as described previously (Krizek, 2003).

### Accession Numbers

ChIP-Seq sequences are available from Sequence Read Archive (https://www.ncbi.nlm.nih.gov/sra) accession number PRJNA685489. Read-only reviewer link is https://dataview.ncbi.nlm.nih.gov/object/PRJNA685489?reviewer=57ngpibcj17sg4sl6v955gpeev Version-controlled source code used to process and analyze data are available from https://bitbucket.org/krizeklab. Sequence alignments and coverage graphs are available for interactive visualization within the Integrated Genome Browser (Nicol *et al.* 2009). To view the data in IGB, readers may download and install the software from https://bioviz.org. Once installed, data sets from the study can be opened within IGB by selecting the latest Arabidopsis thaliana genome and then choosing the ChIP-Seq folder within the Available Data Sets section of the Data Access Panel.

## Acknowledgements

We thank Detlef Weigel for the BJ36_AlcA and pMLBart_AlcR plasmids. This work was supported by National Science Foundation (NSF) grant IOS 1354452. The Integrated Genome Browser software was supported by National Institutes of Health grants R01-GM103463 and R01-GM121927. Data hosting is provided by the SciDas project, funded by NSF award 1659300 to PI Frank (Alex) Feltus.

## Supplemental Data

**Supplemental Figure S1. DNA motifs with similarity to BPC and bHLH binding motifs are over-represented in ANT and AIL6 ChIP-Seq binding peaks.**

**Supplemental Figure S2. ANT and AIL6 bind directly to regulatory regions associated with the floral organ identity gene *PI* but *PI* expression is not altered after downregulation of *AIL6* expression in *35S:ALCR/AlcA:AIL6-amiR ant* inflorescences.**

**Supplemental Figure S3. ANT binds to the *AG* intron *in vitro* and can activate transcription through this binding site in yeast.**

**Supplemental Figure S4. ANT and AIL6 binding peaks within *AP3* and *AG* regulatory regions overlap those of other floral regulators.**

**Supplemental Figure S5. ANT and AIL6 binding peaks within *ANT, AIL6* and *AIL7* regulatory regions overlap those of other floral regulators.**

**Supplemental Figure S6. ANT and AIL6 ChIP-Seq binding peaks in *BB*, *ROT3*, *AN3* and *XTH9* genomic regions.**

**Supplemental Data S1. Genes associated with ANT and AIL6 stage 3 ChIP-Seq binding peaks**

**Supplemental Data S2. GO terms overrepresented in genes associated with ANT stage 3 ChIP-Seq binding peaks**

**Supplemental Data S3. GO terms overrepresented in genes associated with AIL6 stage 3 ChIP-Seq binding peaks**

**Supplemental Data S4. AIL6 peaks that lack an overlapping ANT peak**

**Supplemental Data S5. GO terms overrepresented in genes associated with both ANT and AIL6 stage 3 ChIP-Seq binding peaks**

**Supplemental Data S6. GO terms overrepresented in genes that are differentially expressed in *ant ail6* inflorescences and bound by both ANT and AIL6 in stage 3 flowers**

**Supplemental Data S7. GO terms overrepresented in genes that are differentially expressed in dex treated *35S:ANT-GR* inflorescences and bound by both ANT and AIL6 in stage 3 flowers.**

## Author Contributions

BAK and AEL designed the research. BAK carried out the ChIP-Seq experiments, generated and characterized *35S:ALCR/AlcA:ANT-IR* lines, and performed RT-qPCR, gel mobility shift experiments and yeast one hybrid assays. BAK and JMH performed the ChIP-qPCR. HH constructed the *35S:ALCR/AlcA:AIL6-amiRNA* plasmid and generated *35S:ALCR/AlcA:AIL6-amiRNA* lines, ATB characterized *35S:ALCR/AlcA:AIL6-amiRNA* lines and performed RT-qPCR, and BAK, NF and AEL carried out data analyses. BAK wrote the manuscript. All authors edited the manuscript.

